# Sex-specific perturbations of neuronal development caused by mutations in the autism risk gene *DDX3X*

**DOI:** 10.1101/2024.11.22.624865

**Authors:** Adele Mossa, Lauren Dierdorff, Jeronimo Lukin, Yeaji Park, Chiara Fiorenzani, Zeynep Akpinar, Marta Garcia-Forn, Silvia De Rubeis

## Abstract

*DDX3X* is an X-linked RNA helicases that escapes X chromosome inactivation and is expressed at higher levels in female brains. Mutations in *DDX3X* are associated with intellectual disability (ID) and autism spectrum disorder (ASD) and are predominantly identified in females. Using cellular and mouse models, we show that *Ddx3x* mediates sexual dimorphisms in brain development at a molecular, cellular, and behavioral level. During cortical neuronal development, *Ddx3x* sustains a female-biased signature of enhanced ribosomal biogenesis and mRNA translation. Female neurons display higher levels of ribosomal proteins and larger nucleoli, and these sex dimorphisms are obliterated by *Ddx3x* loss. *Ddx3x* regulates dendritic outgrowth in a sex- and dose-dependent manner in both female and male neurons, and dendritic spine development only in female neurons. Further, ablating *Ddx3x* conditionally in forebrain neurons is sufficient to yield sex-specific changes in developmental outcomes and motor function. Together, these findings pose *Ddx3x* as a mediator of sexual differentiation during neurodevelopment and open new avenues to understand sex differences in health and disease.

## INTRODUCTION

Mutations in the X-linked RNA helicase *DDX3X* are a leading cause of intellectual disability (ID) and autism spectrum disorder (ASD) ^1–6^. While ASD and ID are disproportionally diagnosed and studied in males ^7^, *DDX3X* mutations have a female bias in prevalence, thus providing a unique window into sex differences in neurodevelopment. *DDX3X* mutations have in fact an X-linked semi-dominant pattern of inheritance ^8^. Over 95% of affected individuals are females with *de novo* loss-of-function mutations leading to haploinsufficiency or missense mutations ^2,3,5,9^. The few identified males have missense mutations typically inherited from asymptomatic mothers and thought to act as hypomorphic alleles ^2,3,5,10^.

*DDX3X* encodes an ATP-dependent DEAD/DEAH-box RNA helicase with broad functions in RNA metabolism ^3,11–13^, including regulating the organization of RNA-containing phase-separated organelles ^14^ and the translation of mRNAs with highly structured 5’UTRs ^11,12^. mRNA translation and even the genesis of ribosomes in the nucleoli are finely regulated during brain development ^15–17^. Deficits in these processes have in fact been reported in mouse models of ASD and/or ID ^15,18–21^. The functions of *DDX3X* in the brain are only beginning to emerge ^3,13,22,23^. While *Ddx3x* null mice die in utero ^22,24^, *Ddx3x* haploinsufficient females have delays in meeting postnatal milestones, followed by adult behavioral and neurological deficits ^22^. Evidence in this and other mouse models indicates that *Ddx3x* is indispensable for the development of the hindbrain ^23^ and the forebrain ^3,13,22^. The neocortex is particularly vulnerable, as shown by the congenital brain malformations often diagnosed in individuals with *DDX3X* mutations ^2,3,5^, and the experimental evidence showing that the birth ^3,13^ and laminar subtype specification ^22^ of cortical glutamatergic neurons require *Ddx3x*. In addition to these functions ingrained in prenatal development, prior evidence suggest that *Ddx3x* also regulates neuronal morphogenesis ^25^, which might further compromise the integration of cortical glutamatergic neurons into local circuitry.

*DDX3X* escapes X chromosome inactivation and is expressed from both alleles in human ^26–28^ and mouse ^22,29^ females, including in the brain. *DDX3X* has retained a Y homologous gene (*DDX3Y*), despite only a few ancestral genes having survived on the Y chromosome ^30,31^. For these reasons, *DDX3X* belongs to a class of X-linked genes with unique potential to drive sexual differentiation, through unbalanced expression and/or divergent functions of the X and Y gene partners ^31^. However, the interplay between *DDX3X* and *DDX3Y* remains poorly understood. In brain-conditional mouse models, *Ddx3x* null females have more pronounced brain cytoarchitecture anomalies than *Ddx3x* null males ^13,23^, which also show upregulated *Ddx3y* mRNA ^13,23^, suggesting homeostatic mechanisms of compensation. However, complete ablation of *DDX3X* in male humans and mice is incompatible with life ^5,22,24^. Also, *DDX3Y* deletions cause subfertility/infertility ^32–34^, not brain disorders, in line with the evolution of testis-restricting regulatory regions in primates ^35^. Further, sequence divergence between *DDX3X* and *DDX3Y* yields distinct biochemical properties related to liquid-liquid phase separation and translational regulation ^14^.

All together, these findings place DDX3X at the intersection of X chromosome biology, sexual differentiation, and risk for ID/ASD, and yet no studies have examined the differential effect of *Ddx3x* loss in female and male brains. Here, we aim at addressing this gap by examining the influence of sex, *Ddx3x* gene dosage, and their interaction on neuronal development and behavioral outcomes translationally relevant for *DDX3X* mutations associated with ID and ASD.

## RESULTS

### *Ddx3x* defines the complexity of dendritic arborization in a dosage- and sex-dependent manner

To investigate the influence of sex and *Ddx3x* allelic dosage on neuronal development, we crossed *Ddx3x*^flox/+^ females ^22^ with wild-type males (*Ddx3x*^+/y^) to obtain embryos of four genotypes: control females (*Ddx3x*^+/+^), floxed heterozygous females (*Ddx3x*^+/flox^), control males (*Ddx3x*^+/y^), and floxed males (*Ddx3x*^flox/y^). By crossing *Ddx3x*^flox/+^ females with *Ddx3x*^flox/y^ males we also obtained floxed homozygous females (*Ddx3x*^flox/flox^). Cortical neurons were isolated from embryos at embryonic (E) day 15 across these 5 genotypes and transfected with a bicistronic construct carrying mCherry and CRE (pAAV-Ef1a-mCherry-IRES-Cre) at plating (day in vitro 0, DIV0) (**Fig. 1A**). Neurons were then cultured for 9 days, fixed, and imaged (**Fig. 1A-B**). This genetic manipulation resulted in sparsely labeled mCherry^+^ control female neurons (*Ddx3x*^+/+^), haploinsufficient female neurons (*Ddx3x*^+/−^), null female neurons (*Ddx3x*^−/−^), control male neurons (*Ddx3x*^+/y^), and null male neurons (*Ddx3x*^−/y^) (**Fig. 1A-B**). As intended, mCherry^+^ neurons from female Ddx3x^flox/flox^ embryos or male *Ddx3x*^flox/y^ embryos expressed no DDX3X, while mCherry^+^ neurons from female *Ddx3x*^+/flox^ embryos had depleted DDX3X levels (**Fig. 1B**).

**Figure 1.**
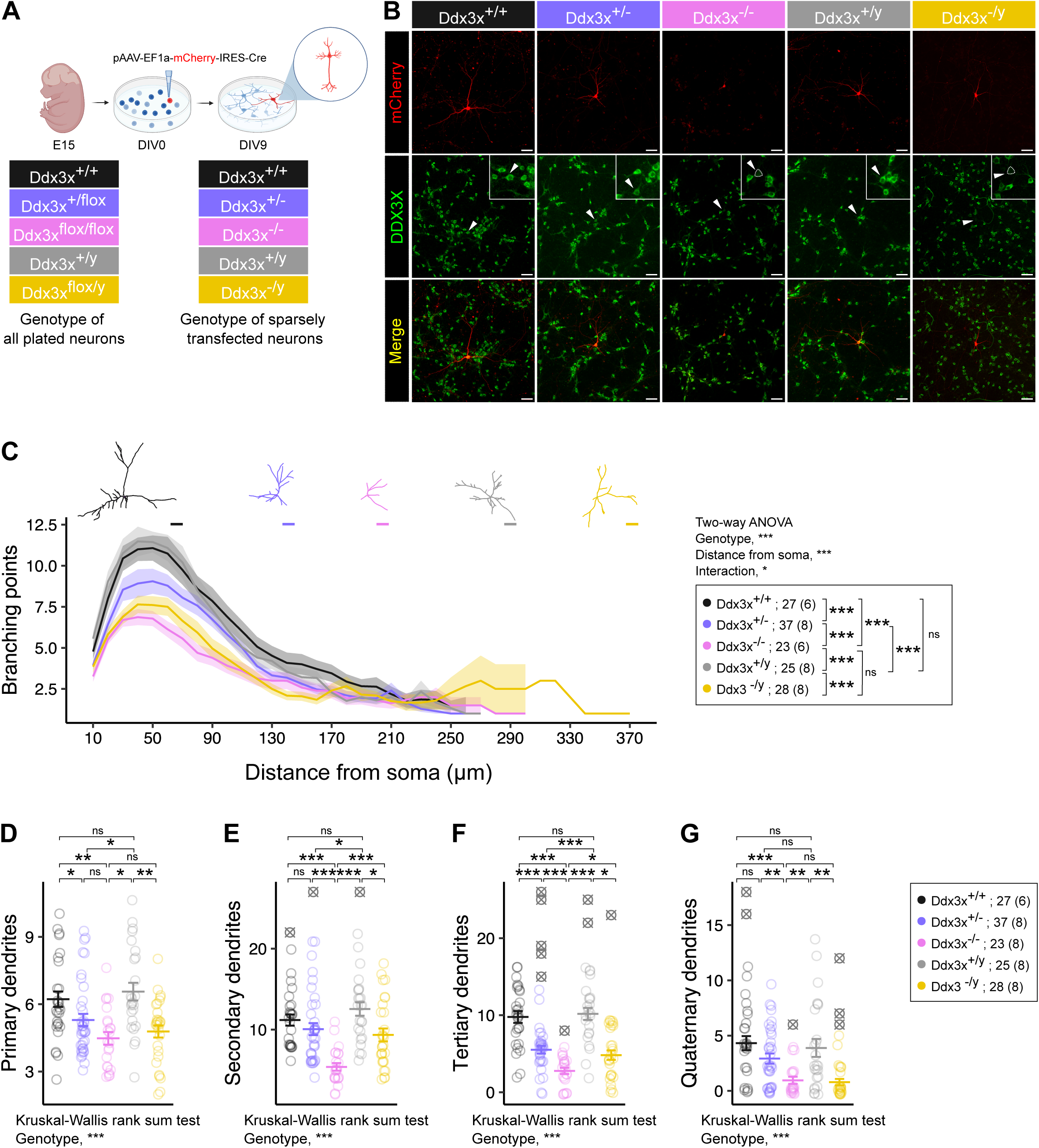
*Ddx3x* regulates dendritogenesis. **A) Experimental design.** Mouse cortical neurons are isolated from embryos at embryonic day (E) 15 from control females (*Ddx3x*^+/+^), flox heterozygous females (*Ddx3x*^+/flox^), flox homozygous females (*Ddx3x*^flox/flox^), control males (*Ddx3x*^+/y^), and flox males (*Ddx3x*^flox/y^). Neurons are transfected the day of plating (day in vitro 0, DIV0) with a bicistronic construct carrying mCherry and CRE, resulting in sparsely labeled control female neurons (*Ddx3x*^+/+^), haploinsufficient female neurons (*Ddx3x*^+/−^), null female neurons (*Ddx3x*^−/−^), control male neurons (*Ddx3x*^+/y^), and null male neurons (*Ddx3x*^−/y^). Analyses are then performed at DIV9. **B) Validation of the *Ddx3x* manipulations in neuronal cultures**. Immunostaining for mCherry (red) and DDX3X (green) on DIV9 cortical neurons, showing the sparse labeling and loss of DDX3X expression in *Ddx3x*^−/y^ and *Ddx3x*^−/−^ null neurons. Scale bar, 50 μm. **C) Loss of *Ddx3x* causes a simplified dendritic arborization.** The plot shows the number of branching points as a function of the distance from the soma in DIV9 neurons across the five genotypes described in panel A and shown in panel B. Representative traces are shown on top of the plot. One-way ANOVA for genotype and distance from the soma, followed by Tukey test with Benjamini-Hochberg correction. **D-G) Loss of *Ddx3x* causes a simplified dendritic arborization.** The plot shows the number of primary (**D**), secondary (**E**), tertiary (**F**) and quaternary dendrites (**G**) on DIV9 cortical neurons across five genotypes described in panel A. Kruskal-Wallis test for genotype, followed by Wilcoxon test with Benjamini-Hochberg correction. In all panels, data were collected blind to genotype and sex; *n* is shown in legend as number of neurons (and number of embryos); mean ± SEM; outliers (shown as ≅), \**P*-value<0.05, \*\**P*-value <0.01, \*\*\**P*-value <0.001.

In line with previous data that siRNA-mediated *Ddx3x* knockdown alters neurite outgrowth ^25^, analysis of DIV9 mCherry^+^ neurons revealed that *Ddx3x* deficiency reduces dendritic complexity: *Ddx3x*^+/−^, *Ddx3x*^−/−^, and *Ddx3x*^−/y^ neurons have simplified dendritic arborizations, as indicated by Sholl analysis measures (**Fig. 1C**) and the number of primary, secondary, tertiary, and quaternary (**Fig. 1D-G**) dendrites.

These cellular deficits are both dosage- and sex-dependent. Female *Ddx3x*^+/−^ and *Ddx3x*^−/−^ neurons differ, indicating a gene dosage effect (**Fig. 1C**, **Fig. 1E-G**). Female *Ddx3x*^+/−^ and male *Ddx3x*^+/y^ neurons both have monoallelic *Ddx3x* expression but show significant differences (**Fig. 1C-F**). Female *Ddx3x*^−/−^ and male *Ddx3x*^−/y^ null neurons showed no overall differences in their dendritic arborization (**Fig. 1C**), but *Ddx3x*^−/−^ neurons had significantly less secondary and tertiary dendrites than *Ddx3x*^−/y^ neurons (**Fig. 1E-F**).

These data indicate that *Ddx3x* is indispensable for the development of a proper dendritic arborization in a cell-autonomous manner in both female and male neurons, and that this function depends on dosage and sex. The results also suggest that *Ddx3y* is unable to fully compensate for *Ddx3x* function in neuronal morphogenesis.

### *Ddx3x* controls the female-biased expression of ribosomal proteins and mRNA translation regulators

We next investigated the molecular changes that accompany these cellular phenotypes. To this end, we infected DIV0 cortical neurons from control females (*Ddx3x*^+/+^), floxed females (*Ddx3x*^+/flox^), control males (*Ddx3x*^+/y^), and floxed males (*Ddx3x*^flox/y^) with AAV8-Ef1a-mCherry-IRES-Cre viral particles. With this strategy, we obtain homogenous populations of *Ddx3x*^+/−^ and *Ddx3x*^−/y^ neurons, and their respective controls. We then performed label-free quantitative proteomics at DIV9, restricting analyses to proteins identified with at least two unique peptides (see Methods), and proceeded to identify up- and down-regulated proteins (indicated in the figure and below with the gene names, **Fig. 2A-C**).

**Figure 2.**
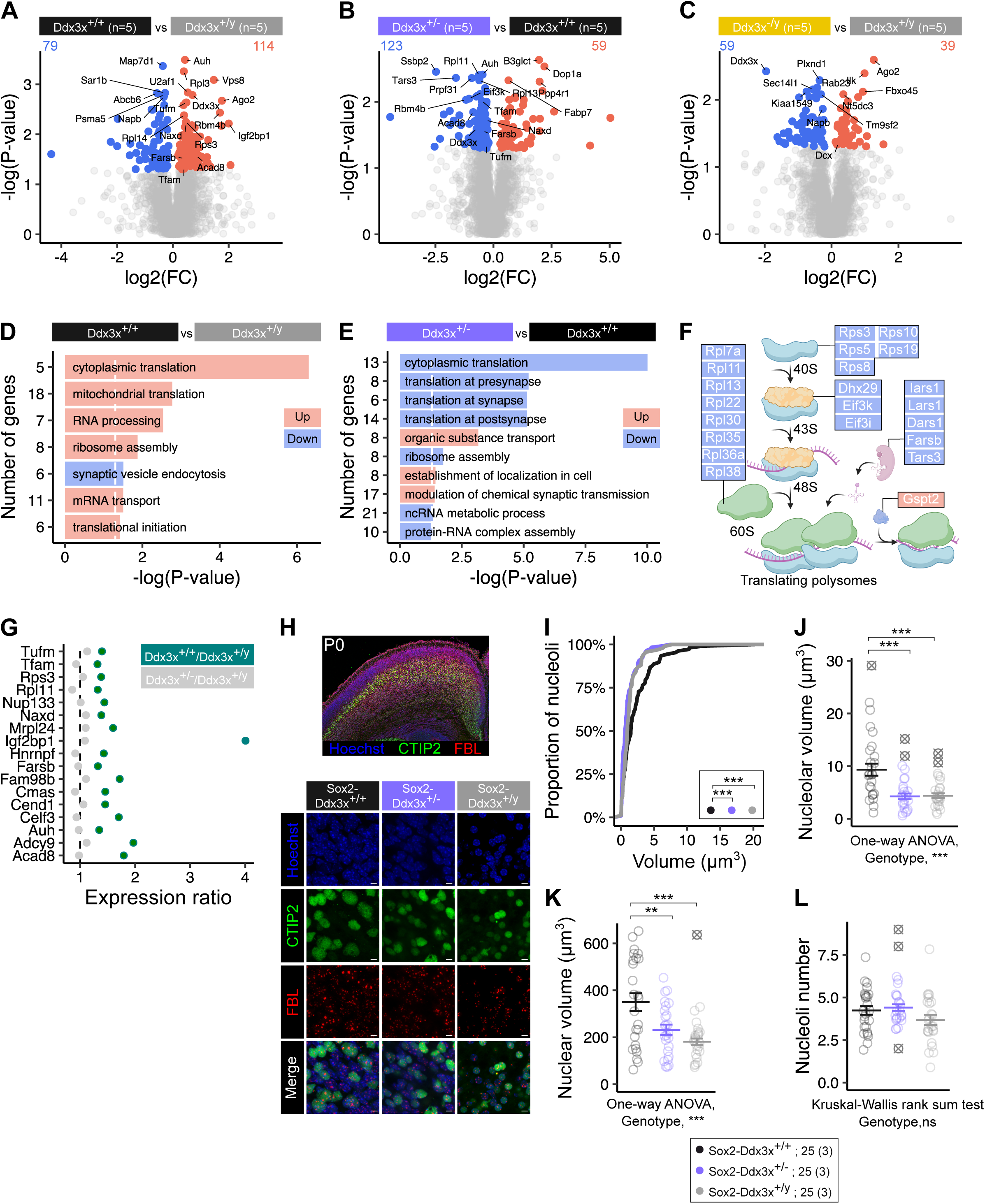
*Ddx3x* regulates the neuronal proteome and mediate a female bias in nucleolar ribosomal biogenesis. **A) Sex influences the neuronal proteome.** Volcano plots of the proteomics data contrasting *Ddx3x*^+/+^ and *Ddx3x*^+/y^ neurons, generated using a viral-mediated transduction approach similar to that delineated in **Fig. 1A**. FC is fold change; grey dotted line denotes *P*-value=0.05. **B) *Ddx3x* haploinsufficiency influences the neuronal proteome.** As in A, contrasting *Ddx3x*^+/+^ and *Ddx3x*^+/−^ neurons. **C) *Ddx3x* ablation in male neurons influences the neuronal proteome.** As in A, contrasting *Ddx3x*^+/y^ and *Ddx3x*^−/y^ neurons. **D) Proteins with increased expression in female neurons converge on RNA processing and mRNA translation.** Enriched Gene Ontology (GO) terms for proteins up- (red) or down-regulated (blue) in *Ddx3x*^+/+^ neurons compared to *Ddx3x*^+/y^ neurons (number of genes in each GO term on the Y axis; white dotted line denotes *P*-value=0.05). **E) Proteins with reduced expression in *Ddx3x* haploinsufficient neurons converge on RNA processing and mRNA translation.** Enriched GO terms for proteins up- (red) or down-regulated (blue) in *Ddx3x*^+/−^ neurons compared to *Ddx3x*^+/+^ neurons, as in panel D. **F) Proteins dysregulated in *Ddx3x* haploinsufficient neurons involved in cytoplasmic mRNA translation.** The scheme summarizes the steps of translational initiation, elongation, and termination. Proteins (indicated by gene names) downregulated in *Ddx3x*^+/−^ neurons compared to *Ddx3x*^+/+^ neurons are shown in blue, while upregulated proteins are shown in red. **G) *Ddx3x* haploinsufficiency obliterates the female bias in expression for 17 proteins.** The plot shows the fold change for the proteins listed on the Y axis by comparing *Ddx3x*^+/+^ compared to *Ddx3x*^+/y^ neurons or *Ddx3x*^+/−^ compared to *Ddx3x*^+/y^ neurons. Proteins are indicated by gene names; grey dotted line denotes reference expression in *Ddx3x*^+/y^ neurons. **H) Sex dimorphisms in nucleoli depend on *Ddx3x*.** Upper panel, representative image of a P0 cortex stained for and Hoechst (blue), CTIP2 (green), Fibrillarin (FBL, red). Lower panel, representative image of magnified regions in the primary motor cortex from *Ddx3x*^+/+^, *Ddx3x*^+/−^, and *Ddx3x*^+/y^ neonates stained for Hoechst (blue), CTIP2 (green), Fibrillarin (FBL, red). **I) Nucleoli are on average larger in female neurons, but not if they have monoallelic expression of *Ddx3x*.** The plot shows the empirical cumulative distribution function of the nucleoli volume across the three genotypes. Legend is shown at the bottom; Kolmogorov–Smirnov test. **J) Total nucleolar volume is higher in female neurons, but not if they have monoallelic expression of *Ddx3x*.** Plot showing the sum volume of all the nucleoli in the nucleus across the three genotypes. Legend is shown at the bottom; One-way ANOVA for genotype, followed by Student’s t test with Benjamini-Hochberg correction. **K) Nuclear volume is higher in female neurons, but not if they have monoallelic expression of *Ddx3x*.** Plot showing the total nuclear volume across the three genotypes. Legend is shown at the bottom; One-way ANOVA for genotype, followed by Student’s t test with Benjamini-Hochberg correction. **L) Number of nucleoli do not differ by genotype.** Plot showing the total number of nucleoli across the three genotypes. Legend is shown at the bottom; Kruskal-Wallis test for genotype, followed by Wilcoxon test with Benjamini-Hochberg correction. In all panels, data were collected blind to genotype and sex; mean ± SEM; outliers (shown as ≅), \**P*-value<0.05, \*\**P*-value <0.01, \*\*\**P*-value <0.001.

When comparing *Ddx3x*^+/+^ and *Ddx3x*^+/y^ genotypes, we found 193 dysregulated proteins (*P*-value<0.05, see **Table S1A**), specifically 114 proteins with higher expression in female neurons than male neurons, and 79 proteins with the opposite sex bias in expression (**Fig. 2A**). *Ddx3x* is one of the most differentially expressed proteins based on sex (**Fig. 2A, Table S1A**), in agreement with our prior data showing higher *Ddx3x* mRNA and protein expression in cortices of female mice compared with male mice ^22^. Interestingly, the only other X-linked protein detected, *Ubl4a* (**Table S1A**), also shows higher expression in female neurons, suggesting that this gene might escape X chromosome inactivation in murine neurons. However, no sex bias in expression is detected for *Ubl4a* in human tissues ^27^ (**Fig. S1A**). Also, the female-biased expression for 4 genes in addition to *Ddx3x* (*Tufm, Scfd2*, *Lrpprc*, and *Eif3i*) is conserved in the human cortex, as shown by bulk transcriptomic data ^28^ (**Fig. S1B**) (see **Supplemental Note**).

When comparing *Ddx3x*^+/−^ and *Ddx3x*^+/+^ genotypes, we found 182 dysregulated proteins (*P*-value<0.05, see **Table S1B**), specifically 59 up-regulated and 123 down-regulated proteins (including *Ddx3x*) (**Fig. 2B**). When comparing *Ddx3x*^−/y^ and *Ddx3x*^+/y^ genotypes, we found 98 dysregulated proteins (*P*-value<0.05, see **Table S1C**), specifically 39 up-regulated and 59 down-regulated proteins (including *Ddx3x*) (**Fig. 2C**). Previous analyses have detected *Ddx3y* mRNA in the brain ^13,23^. However, since *DDX3Y* has been shown to undergo post-transcriptional restriction in male germ cells ^35,36^, we analyzed our proteomics data in search of DDX3Y-specific peptides in male neurons. We identified a DDX3Y-specific peptide (KPILVATAVAAR in DDX3Y, Uniprot #Q62095; SPILVATAVAAR in DDX3X, Uniprot # Q62167) only the *Ddx3x*^−/y^ dataset, corroborating the expression of this protein, albeit at low levels, in male neurons.

We then interrogated biological processes annotated in Gene Ontology (GO) with the datasets of 193 (**Fig. 2A**), 182 (**Fig. 2B**), and 98 (**Fig. 2C**) proteins dysregulated in *Ddx3x*^+/+^, *Ddx3x*^+/−^, *Ddx3x*^−/y^ neurons, respectively. Using Fisher’s exact test with Bonferroni correction, we found that of the 193 proteins with a sex bias in expression, the 114 upregulated in female neurons converge on GO terms related to RNA processing and mRNA translation terms (**Fig. 2D**). Intriguingly, the GO results of the proteins dysregulated in *Ddx3x*^+/−^ when compared with *Ddx3x*^+/+^ neurons offer a specular scenario, with an enrichment of mRNA translation terms for downregulated proteins in *Ddx3x*^+/−^ neurons (**Fig. 2E**). For example, proteins in the top enriched term “cytoplasmic translation” have higher expression in female neurons than in male neurons, but reduced expression in female neurons with *Ddx3x* haploinsufficiency (**Fig. 2D-E**). Amongst the proteins with reduced expression in *Ddx3x*^+/−^ compared to *Ddx3x*^+/+^ neurons and implicated in mRNA translation are 5 tRNA synthetases, 5 ribosomal proteins of the small 40S subunit, 3 initiation factors, 8 ribosomal proteins of the large 60S subunit (**Fig. 2F**), and 5 proteins involved in mitochondrial translation (*Mrps18b*, *Mrpl24*, *Tufm*, *Mrps26*, *Mrps30*). The brain-specific form of the translation termination factor eRF3 ^37^ ERF3B (encoded by *Gspt2*), has enhanced expression (**Fig. 2F**). The observed reduction in ribosomal biogenesis aligns with findings in a *DDX3X* knockdown model in lymphoma cell lines ^38^. Strikingly, 7 of the 13 ribosomal proteins reduced in *Ddx3x^+/−^* neurons are encoded by mRNAs interacting with DDX3X based on iCLIP data in lymphoma ^38^ (i.e., *Rps3*, *Rps5*, *Rps8*, *Rps19*, *Rpl11*, *Rpl13*, *Rpl30*), and 5 were found to have reduced translational efficiency in lymphoma ^38^ (i.e., *Rps3*, *Rps5*, *Rps19*, *Rpl11*, *Rpl13*). These data suggest that in cortical neurons *Ddx3x* regulates the expression of core components of the protein synthesis machinery.

By contrast, we found no significant enrichment for the 98 proteins dysregulated in *Ddx3x*^−/y^ compared with *Ddx3x*^+/y^ neurons. Several of the proteins dysregulated have been shown to be indispensable for proper dendritic development, e.g. *Dcx* ^39^ and *Plxnd1* ^40^, or for neuronal migration, e.g., *Rab23* ^41^ and *Fbxo45* ^42^ (whose human orthologue is part of the genomic interval of the 3q29 Microduplication Syndrome ^43^), suggesting that the similar morphogenesis defects observed in *Ddx3x*^+/−^ and *Ddx3x*^−/y^ neurons (**Fig. 1**) result from distinct molecular mechanisms. Further corroborating divergent molecular processes in female and male *Ddx3x* mutant neurons, the differentially expressed proteins in *Ddx3x*^−/y^ (but not those dysregulated in *Ddx3x*^+/−^ neurons) are enriched for proteins encoded by targets of two RNA-binding proteins associated with ID/ASD, FMRP and RBFOX (**Fig. S1D**).

To gain insights on the disease relevance of the differentially expressed proteins in these three datasets (**Fig. 2A-C**), we examined the intolerance to loss-of-function genetic variation in their human orthologues using the well-established metrics of loss-of-function observed/expected upper bound fraction (LOEUF) ^44^ (**Supplemental Note**). Human orthologues of sex-biased proteins and proteins dysregulated in *Ddx3x* mutant neurons, either female or male, all display skewed distribution toward lower LOEUF deciles (**Fig. S1C**), in keeping with the high genomic constraints for genes expressed in brain. Also, human orthologues of the sex-biased proteins include genes associated with severe neurodevelopmental disorders (DDG2P ^45^ genes), ASD risk genes (ASC ^46^ genes), and/or epilepsy (EPI ^47–49^ genes), albeit with no statistically significant enrichment (**Supplemental Note**). For example, 18 proteins encoded by DDG2P genes have higher expression in female neurons (e.g., *Ddx3x*, *Auh*, *Tufm*) and another 8 have higher expression in male neurons (e.g., *Napb*, *Shank2*, *Ank2*) (**Table S1**). 5 proteins encoded by ASC gene orthologues have a sex bias in expression, and they all have higher expression in male neurons (*Adcy5*, *Coro1a*, *Ank2*, *Shank2*, *Cul3*) (**Table S1**). 30 proteins encoded by DDG2P gene orthologues have altered expression in *Ddx3x*^+/−^ neurons, specifically 10 up-regulated proteins (e.g., *Shank2*, *Nrxn3*) and 20 down-regulated proteins (e.g., *Auh*, *Ccdc88a*, *Tufm*). 13 proteins encoded by DDG2P gene orthologues have altered expression in *Ddx3x*^−/y^ neurons, specifically 7 up-regulated proteins (e.g., *Dcx*, *Adsl*, *Shank1*) and 6 down-regulated proteins (e.g., *Rab23*, *Napb*).

These data map the molecular correlates of the deficits in neuronal morphogenesis caused by perturbations in *Ddx3x* (**Fig. 1**) and show that there are sex-specific *Ddx3x* signatures, including alterations of ribosomal proteins and translational regulators exclusively in female neurons.

### *Ddx3x* mediates sex dimorphisms in neuronal ribosomal biogenesis

Female neurons have higher dosage requirements for *Ddx3x* (e.g., **Fig. 2A**, **Fig. S1A-B**), as it has evolved as an X chromosome escapee gene ^22,27,29^, and display enhanced expression of components of the protein synthesis machinery (**Fig. 2D**). *Ddx3x^+/−^* on the contrary have dampened expression of ribosomal proteins (**Fig. 2E-F**). Therefore, we conjectured that *Ddx3x* might mediate the female-biased surplus of protein synthesis. To test this hypothesis, we verified if the higher expression of proteins in female neurons requires *Ddx3x*. We started by intersecting proteins with altered levels in *Ddx3x*^+/−^ neurons (**Fig. 2B**) with those showing differential expression based on sex (**Fig. 2A**) and identified 37 proteins (**Table S2**). We then filtered for proteins that have at least 30% higher expression in female neurons compared to male neurons (Expression*_Ddx3x_*_+/+_/Expression*_Ddx3x_*_+/y_>1.3) and identified 20 proteins. Of these 20, 17 have similar levels of expression when comparing female haploinsufficient neurons and male control neurons (Expression*_Ddx3x_*_+/-_/Expression*_Ddx3x_*_+/y_=1±0.15), indicating that *Ddx3x* deficiency obliterates their female bias in expression (**Fig. 2G**, **Table S2**). 6 are encoded by genes whose human orthologues are DDG2P genes (*TUFM*, *TFAM*, *NUP133*, *NAXD*, *FARSB*, and *AUH*) and 1 (*TFUM*) also shows significantly higher expression in human cortices from female donors than male donors (**Fig. S1B**). The remaining 3 proteins (*Rbm4b*, *Pds5b*, *Ssbp1*), do not meet these stringent criteria but still show an attenuation of their sex bias (**Table S2**).

Of these 17 proteins, 5 are implicated in mRNA translation (*Tufm*, *Rps3*, *Rpl11*, *Mrpl24*, *Farsb*; **Fig. 2F-G**) and another 4 are RNA-binding proteins (*Igf2bp1*, *Hnrnpf*, *Celf3*, *Auh*). *Rps3* and *Rpl11* were also detected in a DDX3X iCLIP dataset ^38^. These data suggest that female neurons have enhanced protein synthesis machinery, including ribosome biogenesis, and that this female bias depends on *Ddx3x*. Ribosome biogenesis takes place in the nucleolus, a phase-separated organelle in the nucleus. *DDX3X* ^14^, as well as other DEAD-box RNA helicases ^50^, have proven to be critical for liquid-liquid phase separation. Therefore, we conjectured that the *Ddx3x*-dependent signature in female neurons might reflect changes in nucleolar dynamics. To directly test this hypothesis, we sought to examine the number and volume of nucleoli *ex vivo* and in culture. We took advantage of the *Sox2*-*Ddx3x*^+/−^ mouse model we have previously generated and characterized ^22^. While *Sox2-Ddx3x*^−/y^ die *in utero*, *Sox2*-*Ddx3x*^+/−^ haploinsufficient females show cellular, behavioral, and neurological alterations reminiscent of clinical phenotypes of *DDX3X* mutations ^22^. Building on our previous observation of alterations of glutamatergic cortical neurons expressing the transcription factor CTIP2 in *Sox2*-*Ddx3x*^+/−^ females ^22^, we examined nucleoli in CTIP2+ neurons of the motor cortex of control female (*Sox2-Ddx3x*^+/+^), haploinsufficient female (*Sox2*-*Ddx3x*^+/−^), and control male (*Ddx3x*^+/y^) neonates (**Fig. 2H**). We found that CTIP2+ female neurons have on average larger nucleoli (**Fig. 2I**) and higher total nucleolar volume (**Fig. 2J**), without changes in the number of nucleoli (**Fig. 2L**), when compared to male neurons. We also found that female neurons have larger nuclei than male and female haploinsufficient neurons (**Fig. 2K**). In transfected cells, we found similar trends for total nucleolar volume (**Fig. S2A-B**) and *Ddx3x*-dependent female bias in the number of nucleoli (**Fig. S2A, B**). Importantly, although *Ddx3x*^+/−^ and *Ddx3x*^−/y^ neurons have similar underdeveloped dendrites (**Fig. 1**), *Ddx3x*^−/y^ neurons do not show changes in nucleoli compared with male controls (**Fig. S2**).

These data show that *Ddx3x*^+/−^ female neurons are indistinguishable from male neurons in the expression of proteins with skewed expression in female neurons (**Fig. 2G**) and nucleoli dynamics (**Fig. 2H-L, Fig. S2**), indicating that biallelic expression of *Ddx3x* in female neurons mediates these sex dimorphisms.

### *Ddx3x* regulates the development of dendritic spines in a sex-dependent manner

Following the previous reasoning, we sought to discriminate the effect of sex and *Ddx3x* dosage on the formation and maturation of dendritic spines, given that disruption of dendritic development and/or synaptic development have important repercussions on neural circuits underlying behaviors and are common in mouse models of ASD and related neurodevelopmental disorders ^51–56^. We adopted the same Cre-recombination strategy described above. To isolate the discrete function of *Ddx3x* in this process, sparse transfections were performed at DIV9 and neurons examined at DIV14 (**Fig. 3A**). This way, dendritogenesis proceeds unperturbed until the onset of synaptogenesis, and thus synaptic phenotypes are not confounded by earlier defects in dendritic development (**Fig. 1**). 10 μm long segments at least 30 μm distant from the soma were then examined in DIV14 mCherry^+^ control female neurons (*Ddx3x*^+/+^), haploinsufficient female neurons (*Ddx3x*^+/−^), null female neurons (*Ddx3x*^−/−^), control male neurons (*Ddx3x*^+/y^), and null male neurons (*Ddx3x*^−/y^) (**Fig. 3A**). The dendritic segments were examined across primary, secondary, and tertiary dendrites.

**Figure 3.**
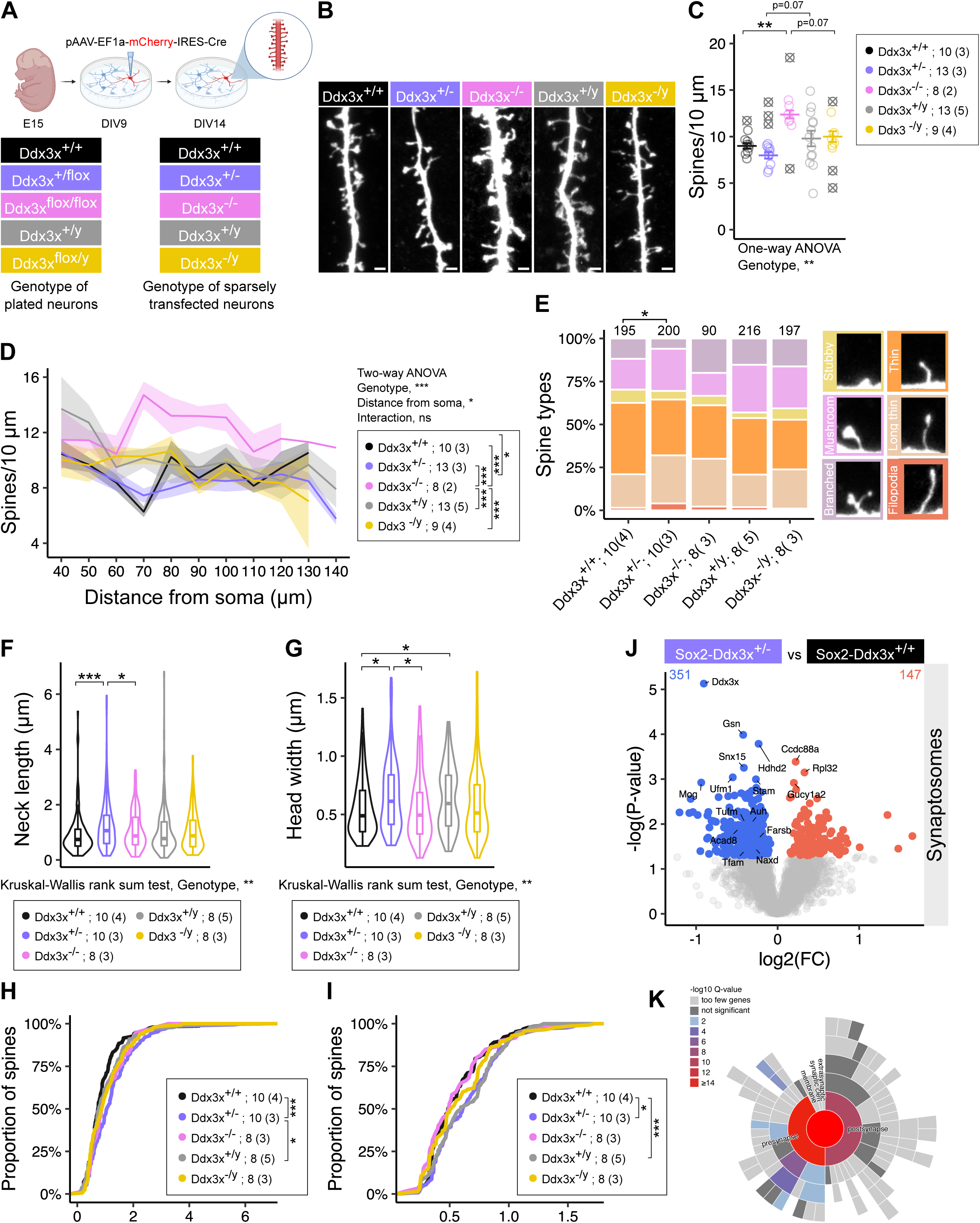
*Ddx3x* regulates the formation and development of dendritic spines. **A) Experimental design.** Mouse cortical neurons are isolated from *Ddx3x*^+/+^, *Ddx3x*^+/flox^, *Ddx3x*^flox/flox^, *Ddx3x*^+/y^, and *Ddx3x*^flox/y^ E15 embryos. At DIV9, neurons are transfected with the construct carrying mCherry and CRE, resulting in sparsely labeled *Ddx3x*^+/+^, *Ddx3x*^+/−^, *Ddx3x*^−/−^, *Ddx3x*^+/y^, and *Ddx3x*^−/y^ neurons. Analyses are performed at DIV14. **B) *Ddx3x* dosage influences dendritic spine density.** Confocal images of 10 μm dendritic segments 50 μm distant from the soma, showing mCherry-expressing dendritic shafts and spines across the five genotypes. Scale bar, 1 μm. **C) *Ddx3x* loss in female (but not male) neurons alters spine density.** The plot shows the number of dendritic spines in 10 μm across the five genotypes. One-way ANOVA for genotype, followed by Student’s t test with Benjamini-Hochberg correction. **D) Dendritic density varies as a function of the distance from the soma.** The plot shows the number of dendritic spines in 10 μm as a function of distance from soma (values on the X axis represent the upper boundary of the segment, e.g., 50 μm marks the 40-50 μm segment). Two-way ANOVA for genotype and distance from the soma, followed by Tukey test with Benjamini-Hochberg correction. **E) *Ddx3x* haploinsufficiency alters the balance of spine subtypes.** The plot shows the proportion of dendritic spines across the six spine subtypes, defined as indicated in the Materials & Methods section (*n* spines above each bar). Chi-square test for genotype. **F-G) *Ddx3x* haploinsufficiency results in increased spine neck length,** as shown by the average spine neck length (E) and the empirical cumulative distribution function of the spine neck length measures (F). Panel E, Kruskal-Wallis test for genotype, followed by Wilcoxon test with Benjamini-Hochberg correction. Panel F, Kolmogorov–Smirnov test. **H-I) *Ddx3x* haploinsufficiency results in increased spine head width,** as in E-F but for the spine head width. **J-K) *Ddx3x* haploinsufficiency changes the synaptic proteome.** Volcano plot (J) and SynGO ^58^ enrichment plot (K) of the quantitative proteomics data on synaptosomes isolated from P21 *Sox2-Ddx3x*^+/+^ (purple) and *Sox2-Ddx3x*^+/−^ (light purple) mice. FC is fold change; grey dotted line denotes *P*-value=0.05. In all panels, data were collected blind to genotype and sex; *n* is shown in legend as number of neurons (and number of embryos); mean ± SEM; outliers (shown as ≅), \**P*-value<0.05, \*\**P*-value<0.01, \*\*\**P*-value<0.001.

Quantifications of dendritic density in DIV14 mCherry^+^ neurons revealed that intact *Ddx3x* expression is indispensable for spine formation in females but not in males: *Ddx3x*^−/−^, but not *Ddx3x*^+/−^ or *Ddx3x*^−/y^ neurons, have more dendritic spines than control neurons (**Fig. 3B-C**). When measuring spine density as a function of distance from the soma (**Fig. 3D**), we noted distinguishable patterns for male or female neurons (e.g., at 60-70 μm distance from the soma), except for *Ddx3x*^−/−^ neurons. We also observed that the increased number of spines in *Ddx3x*^−/−^ neurons is most prominent after 60 μm distance from the soma (**Fig. 3D**).

To further examine spine morphological characteristics, we measured the neck length and the head width of individual dendritic spines. *Ddx3x*^+/−^ female neurons displayed increased neck length (**Fig. 3F**) and a shifted curve for the empirical distribution frequency of this measure (**Fig. 3H**), when compared to *Ddx3x*^+/+^ or *Ddx3x*^−/−^ female neurons. *Ddx3x*^+/−^ female neurons also showed increased head width (**Fig. 3G**) and a shifted curve of empirical distribution frequency (**Fig. 3I**), when contrasted to *Ddx3x*^+/+^ or *Ddx3x*^−/−^ female neurons. We then used these measures to classify spines into 6 morphological types: branched, mushroom, stubby, thin, long thin, and filopodia (**Fig. 3E**) (see Methods). When considering spines cumulatively, we found that only *Ddx3x*^+/−^ female neurons have a distinct profile, with a surplus of mushroom-like and long thin spines at the expense of thin spines (**Fig. 3E**). Collectively, these data are in contrast with prior observations of reduced spine density, excess of mushroom-like spines, and increase of filopodia after acute knockdown of *Ddx3x*, again without considering the variable of sex ^25^.

To explore the molecular correlates of the altered synaptic development of the *Ddx3x*^+/−^ neurons, we delineated the proteomic profile of synapses purified from *Sox2*-*Ddx3x*^+/−^ females at postnatal day 21, a peak for synaptogenesis. We isolated cortical synaptosomes from *Sox2*-*Ddx3x*^+/−^ and control *Sox2*-*Ddx3x*^+/+^ mice and performed label-free quantitative proteomics. As for prior experiments, we restricted analyses to proteins identified with at least two unique peptides (see Methods) and proceeded to identify up- and down-regulated proteins (indicated in the figure and below with the gene names, **Fig. 3J**).

When comparing *Ddx3x*^+/+^ and *Ddx3x*^+/−^ synaptosomes, we found 498 dysregulated proteins (*P*-value<0.05, **Table S3**), specifically 147 proteins up-regulated and 351 down-regulated in *Ddx3x*^+/−^ synaptosomes (**Fig. 3J**). *Ddx3x* was also found to be expressed at synapses in line with a prior synaptic proteomic dataset ^57^ and, as expected, reduced in expression in *Ddx3x*^+/−^ synaptosomes (**Fig. 3J, Table S3**). As expected, the differentially expressed proteins are enriched in synaptic terms (**Fig. 3K**), as 114 of the 346 proteins dysregulated in *Ddx3x*^+/−^ synaptosomes are *bona fide* synaptic proteins as defined by the SynGO consortium ^58^. We then crossed the list of genes encoding the 498 differentially expressed proteins with the DDG2P genes ^45^, ASC genes ^46^, and EPI genes ^47–49^ (see **Supplemental Note**), and found a statistically significant enrichment for DDG2P genes (**Fig. S1D**). 84 are encoded by DDG2P genes, 9 by ASC genes (4 of which also captured by DDG2P; *Psmd12*, *Nrxn1*, *Scn2a*, and *Plxna1*), and 3 by EPI genes (all 3 captured by DDG2P genes, *Ube3a*, *Eef1a2*, and *Scn2a*). At the intersection of the differential expressed proteins, established SynGO genes, and disease risk genes are critical regulators of synaptic function, including *Dlg3*, *Nrxn1*, *Nrxn2*, *Scn2a*, *Cntnap2* and *Rac1*.

When crossing the list of dysregulated proteins in *Ddx3x^+/−^*neurons (**Fig. 2B**) and synaptosomes (**Fig. 3J**), we and found 19 shared proteins in addition to *Ddx3x*, including *Rpl22a* and *Ccdc88a* (also known as Girdin). Amongst these 19, there are 6 that are also expressed at higher levels in female neurons in a *Ddx3x*-dependent manner (*Auh*, *Tufm*, *Naxd*, *Acad8*, *Farsb*, and *Tfam*) (**Fig. 2G**).

Our data indicate that *Ddx3x* influences the development of dendritic spines in a cell-autonomous manner. This regulation is female-specific, as *Ddx3x* null male neurons are unaffected. The effect of dosage in female neurons is complex, as haploinsufficiency affects spine morphology and the synaptic proteome but not density, and complete loss affects density but not spine morphology.

### Loss of *Ddx3x* in the forebrain leads to developmental delays and adult motor deficits

We then next sought to assess whether the interaction between *Ddx3x* dosage and sex observed at the molecular (**Fig. 2**) and cellular level (**Fig. 1, 3**) translates to behavioral outcomes.

Complete knockout of *Ddx3x* is incompatible with life in male mice ^22,24^, while *Ddx3x^+/−^*females have postnatal motor delays, adult motor deficits, hyperactivity, anxiety-related behaviors, and cognitive impairments ^22^. To be able to compare female and male mutant mice, we selected a forebrain-conditional *Ddx3x* knock-out model, based on clinical observations ^2,3,5^ and evidence in mouse models ^3,13,22^ (including those presented here) showing that forebrain neurons are particularly vulnerable to *DDX3X* mutations. To this end, we crossed the *Ddx3x^flox^* line with an *Emx1^IRES^*^−*Cre*^ driver line that expresses Cre under the endogenous Emx1 promoter in the telencephalon as early as E10.5 and in ∼88% of neocortical and hippocampal neurons in the adult brain ^59,60^. Unlike in other Emx1-Cre lines, in Emx1^IRES-*Cre*^ mice EMX1 expression is unaffected ^60^, avoiding a major confounder. We obtained four genotypes: *Emx1-Ddx3x^+/+^* control females, *Emx1-Ddx3x^+/−^* haploinsufficient females, *Emx1-Ddx3x^+/y^* control males, and *Emx1-Ddx3x^−/y^* null males. There was no statistically significant difference between the expected mendelian ratio and the percentages obtained for each genotype over 15 litters. Moreover, *Emx1-Ddx3x^−/y^* mutants did not show reduced viability from birth to the postnatal day 21 and survived to adulthood.

As intended, *Emx1-Ddx3x^−/y^* null males show ablation of DDX3X protein in the forebrain, but not in the midbrain or hindbrain (**Fig. 4A**). *Emx1-Ddx3x^−/y^* null males show delayed growth over the first four weeks of life (**Fig. 4B**), but then normalize in adulthood (**Fig. 4C**). *Emx1-Ddx3x^−/y^* null males show no major physical (**Fig. S3**), sensory (**Fig. S4**), or motor (**Fig. S5**) developmental delays, but have a statistically significant delay in the eruption of the top tooth (**Fig. S3D**) and the acquisition of motor abilities such as grip strength (**Fig. S5E**) and air righting (**Fig. S5F**) show a distinct trajectory over time. The developmental trajectory of the *Emx1-Ddx3x^+/−^*heterozygous females is mostly indistinguishable from that of their control littermates (**Fig. 4B, Fig. S2-5**), except for a sensory delay in the forelimb grasp reflex (**Fig. S4B**). Further, both *Emx1-Ddx3x^−/y^*and *Emx1-Ddx3x^+/−^* juvenile mice display subtle changes suggestive of a wider gait, appearing at P22 in *Emx1-Ddx3x^+/−^* and only at P30 in *Emx1-Ddx3x^−/y^* males (**Fig. S6**). This is in contrast with what we previously observed in *Sox2-Ddx3x^+/−^*mice, which showed a reduction in horizontal and diagonal parameters at P22, resolved by P30 ^22^. Collectively, these data indicate that the previously reported developmental delays observed in the global *Sox2-Ddx3x* model are not exclusively dependent on the forebrain, and that nevertheless there is an interaction between sex and genotype and/or a complex dosage effect, as some phenotypes are detected in haploinsufficient females but not in null males (**Fig. S4B**, **Fig. S6**).

**Figure 4.**
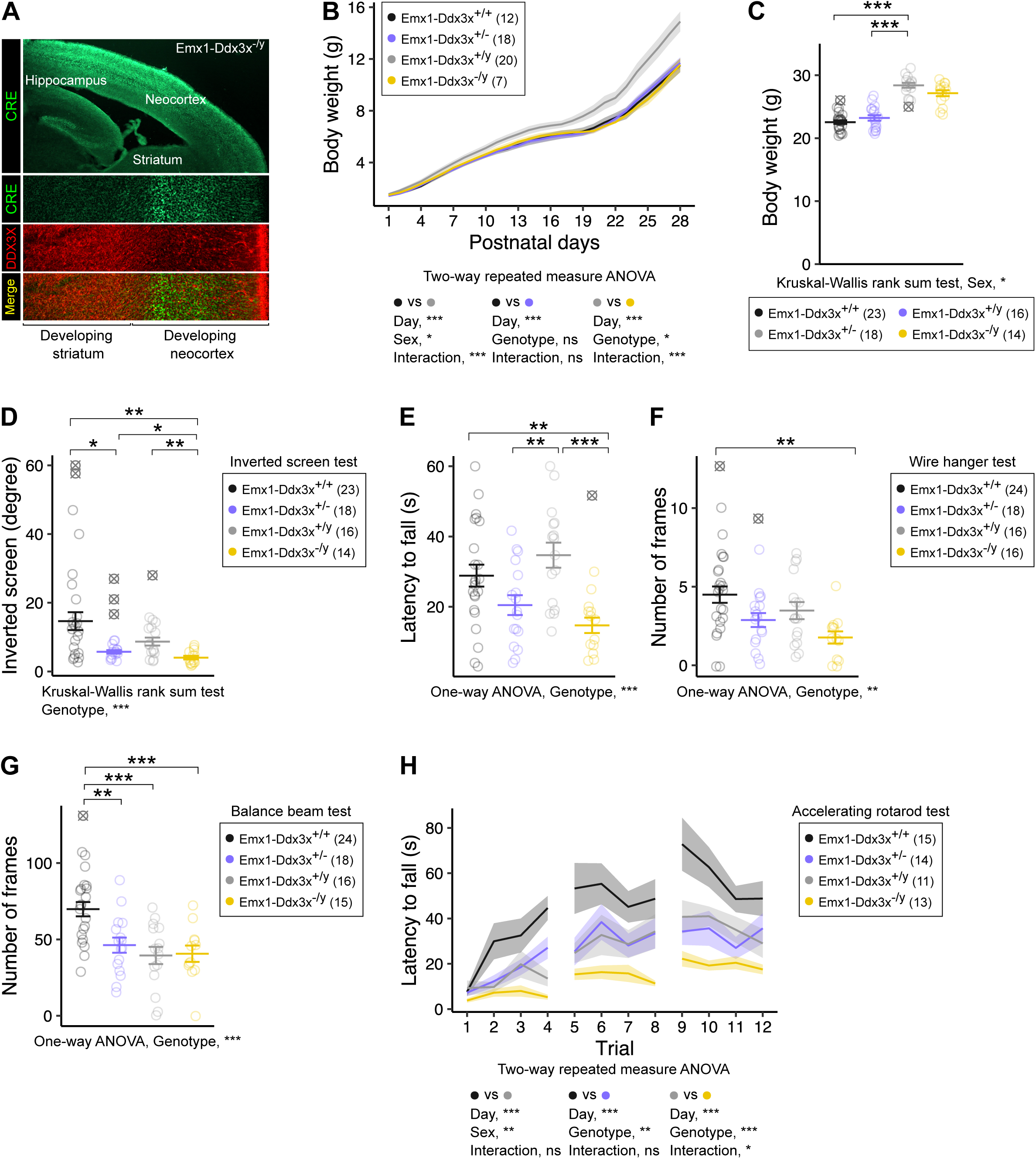
*Ddx3x* loss in the forebrain causes adult motor deficits. **A) Validation of the *Emx1-Ddx3x* knock-out mouse line.** Confocal images of a E18.5 section from an *Emx1-Ddx3x^−/y^* null mouse, showing CRE (green) exclusively in the forebrain, with corresponding loss of DDX3X (red). **B) *Emx1-Ddx3x* null pups have delayed postnatal growth.** Growth curves for *Emx1-Ddx3x^+/+^* control females, *Emx1-Ddx3x^+/−^* haploinsufficient females, *Emx1-Ddx3x^+/y^* control males, and *Emx1-Ddx3x^−/y^* null males. Two-way repeated measure ANOVA for the comparisons shown below the plot. **C) *Emx1-Ddx3x* null males no longer display lower body weight as adults.** Body weight in adult mice. Kruskal–Wallis rank sum test, followed by Wilcoxon signed-rank test with Benjamini-Hochberg correction. **D) *Ddx3x* loss causes reduced grip in inverted screen test,** when compared with *Emx1*-*Ddx3x*^+/+^ female and *Emx1*-*Ddx3x*^+/y^ male controls. Kruskal-Wallis test for genotype, followed by Wilcoxon test with Benjamini-Hochberg correction. **E-F) *Ddx3x* loss reduces endurance in wire hanger test**, as measured by latency to fall (E) and number of frames (F). One-way ANOVA, followed by Student’s t test with Benjamini-Hochberg correction. **G) Sex and *Ddx3x* dosage in the forebrain affect performance on balance beam test**, measured as number of frames walked. One-way ANOVA, followed by Student’s t test with Benjamini-Hochberg correction. **H) Sex and *Ddx3x* dosage in the forebrain affect performance on an accelerating rotarod**, measured as latency to fall. Two-way repeated measure ANOVA for the comparisons shown below the plot. In all panels, data were collected blind to genotype and sex; *n* is shown in legend as number of mice; mean ± SEM; outliers (shown as ≅), \**P*-value <0.05, \*\**P*-value<0.01, \*\*\**P*-value<0.001.

Amongst the most clinically salient phenotypes observed in the *Sox2-Ddx3x* mouse model are motor delays and poor motor coordination and balance ^22^, reminiscent of the pervasive motor manifestations in individuals with *DDX3X* mutations ^2,3,5^. Therefore, we assessed skilled motor functions in the *Emx-Ddx3x* mutant line. During handling one week prior to behavioral testing, we noted no differences in physical appearance or weight (**Fig. 4C**). In terms of baseline activity, we observed reduced inverted screen performance for both *Emx1-Ddx3x^+/−^* and *Emx1-Ddx3x^−/y^* mutants compared to controls (**Fig. 4D**), and reduced number of jumps during spontaneous activity monitoring for *Emx1-Ddx3x^+/−^* females (**Fig. S7B**), but no other changes (**Fig. S7A, S7C-E**).

To assess neuromuscular strength, we employed a wire hanging test (**Fig. 4E-F**). We detected a significant effect of genotype on performance, with *Emx1-Ddx3x^−/y^* null males showing reduced latency to fall from the wire when compared to male littermates (**Fig. 4E**), similar to the reduced endurance observed in *Sox2-Ddx3x^+/−^* mice ^22^. To assess motor coordination and balance, we used a balance beam test and a vertical pole test. When walking on the balance beam, control females covered a larger distance on the beam (**Fig. 4G**) than male littermates, plausibly because of their lower body weight (**Fig. 4C**). In line with observations in *Sox2-Ddx3x^+/−^*mice ^22^, *Emx1-Ddx3x^+/−^* covered a shorter distance on the beam than control female littermates (**Fig. 4G**). No differences in the number of slips were noted (**Fig. S8A**). No sex or genotype differences were observed when mice were climbing down a vertical pole (**Fig. S8B-C**), in contrast to previous findings of impairments in this motor task in *Sox2-Ddx3x^+/−^* mice ^22^. To further test motor coordination and endurance, and extend on motor learning, we used the accelerating rotarod test, with a design of four trials each day, over three days. In line with the outcomes of the balance beam test (**Fig. 4G**), control females outperformed male littermates (**Fig. 4H**). While *Emx1-Ddx3x^+/−^* females were able to learn the task, they showed reduced performance compared to control females (**Fig. 4H**). *Emx1-Ddx3x^−/y^*males showed both impaired motor learning and coordination or endurance when compared to control males (**Fig. 4H**).

These observations indicate that *Ddx3x* deficiency in the forebrain is sufficient to produce motor sequelae reminiscent of those found in the patient population ^61^ and reaffirm the complex sex-by-dosage interactions for the *Ddx3x* locus.

## DISCUSSION

By virtue of its nature as an X chromosome inactivation escapee ^62^, *DDX3X* is one of the top 15 female-biased genes in the human cortex ^28^ (**Fig. S1**), and one of the top 5 proteins with higher expression in female cortical murine neurons (**Fig. 2A**, **Table S1**). In humans, despite the dramatic Y chromosome decay during evolution, *DDX3X* and another 18 escapee genes have retained Y homologues ^30^, suggesting that the equilibrium between the X and Y partners (or the lack thereof) contributes to shape sex dimorphisms ^31^.

Our study provides evidence that *Ddx3x* acts as a mediator of sexual differentiation during brain development. First, we show that *Ddx3x* is necessary for the expression of a female-specific molecular and cellular signature associated with ribosome biogenesis and mRNA translation: female neurons have larger nucleoli and enhanced expression of ribosomal proteins and translational regulators than male neurons, but not if they lack one *Ddx3x* allele (**Fig. 2**). *DDX3Y* has stronger propensity for liquid-liquid phase separation than *DDX3X* ^14^, which might contribute to the lack of nucleolar alterations in *Ddx3x* null male neurons (**Fig. S2**). *Ddx3x* loss in male neurons, in fact, results in a distinct and narrowed impact on proteins directly necessary for neurite outgrowth (**Fig. 2**), with no noticeable nucleolar alterations (**Fig. S2**). Second, *Ddx3x* is indispensable for the formation and maturation of dendritic spines in female neurons only, as spines of male control or null neurons are indistinguishable. Third, female mice outperform males in two motor tasks, but this sex advantage is lost in *Ddx3x* mutant females (**Fig. 2G-H**).

Our findings have also direct relevance for our understanding of how neurodevelopment goes awry when *DDX3X* is mutated. Our observations add to the growing data in the patient population ^2,3,5^ and mouse models ^3,13,22^ indicating that forebrain neurons, particularly in the cortex, are exquisitely vulnerable to *DDX3X* mutations. ∼80% of individuals with a diagnosis of DDX3X syndrome show brain abnormalities suggestive of defective cortical development, including microcephaly, hypoplasia of the corpus callosum, and polymicrogyria ^2,3,5^. Consistently, *Ddx3x* haploinsufficient mice have a thinner cortex with altered cytoarchitecture ^22^. *Ddx3x* conditional loss in the forebrain disrupts cortical neurogenesis and results in microcephaly, as *Emx1-Ddx3x* mutant embryos show an excess of neural progenitors to the expenses of neuronal differentiation ^13^. Here we show that *Ddx3x* expression in the forebrain is also required for intact motor development and function, as both *Emx1-Ddx3x* haploinsufficient female and null male mice have motor deficits, although with significant sex differences (**Fig. 4**).

Our observations add a layer of complexity to the molecular alterations resulting from *DDX3X* mutations. By virtue of its RNA helicase properties and based on data in human cell lines ^11,12^, DDX3X has been postulated to act at the step of translational initiation and facilitate the translation of a subset of mRNAs with 5’UTRs bearing complex secondary structures. When separating by sex, we found two distinct signatures in female and male mutant neurons. In *Ddx3x* female haploinsufficient neurons, we found a general downregulation of constitutive components of the protein synthesis machinery (**Fig. 2**), in line with a previous report in *Ddx3x* mutant lymphoma cells ^38^. The reduced levels of ribosomal factors are accompanied by enhanced translational termination, potentially as a homeostatic response to enhance ribosomes recycling ^63^. These observations, orthogonally corroborated by reduced nucleoli capacity (**Fig. 2**), suggest a general hypoactivity of the ribosomal machinery, that might in turn affect protein synthesis required to sustain dendritic growth, yielding the reduced dendritic branching resulting from *Ddx3x* perturbations (**Fig. 1**). Coherently, genetic mutations in core translational factors dysregulated in *Ddx3x* mutant neurons and/or synapses (*IARS1* ^64^*, DARS* ^65^, *FARSB* ^66^, and *GSPT2* ^67,68^) have been associated with adverse neurodevelopmental outcomes. Further, our observation of nucleolar alterations (**Fig. 2**, **Fig. S2**) adds on an emerging class of genetic conditions associated with nucleolar alterations and defective ribosomal biogenesis ^21,69,70^, including a recent report in a Fragile X syndrome mouse model ^21^.

Our data also show that *Ddx3x* is needed for the development of cortical neurons post-mitotically (**Fig. 1-3**), in line with a previous report ^25^, thus expanding the window of risk for *DDX3X* mutations beyond prenatal development ^3,13^. Synaptic function, in particular, has been pinpointed as a major nexus of risk for ASD ^71^, and genetically engineered mouse models for ASD have also shown deficits in synaptic development ^51,52,72,73^. Importantly, dendritic growth, *de novo* synapse formation, and synapse maturation continue to be plastically reshaped during postnatal periods, including in response to experience ^74–76^, thus offering a more feasible window for therapeutic intervention. Our previous data ^22^ lend support for potential benefits of intervening postnatally on *Ddx3x* mutant mice. In fact, we found that one-year old *Sox2-Ddx3x*^+/−^ mice have more profound motor deficits than naïve 4-month-old *Sox2-Ddx3x*^+/−^ mice, but this motor decline is no longer seen in one-year old mice pre-exposed to behavioral training at 4 months of age. These data suggest that there might be mechanisms maintaining the pathological status, perhaps through the synaptic cellular and molecular alterations we found (**Fig. 1-3**), that could be manipulated postnatally. This might add to the growing body of evidence showing that restoring expression/function of some ASD risk genes in adult mice can alleviate their molecular and/or behavioral deficits ^77–81^. Future studies will be critical to understand how these molecular and cellular alterations can be targeted for the development of novel therapeutics.

## METHODS

### Mouse lines

All animal procedures were approved by the Institutional Animal Care and Use Committee of the Icahn School of Medicine at Mount Sinai. To generate the neuronal cultures for the dendritic and synaptic analyses, we crossed *Ddx3x^flox/+^* females ^22^ with wild-type *Ddx3x*^+/y^ males or *Ddx3x*^flox/y^ males. To generate the *Sox2-Ddx3x* mouse line used for the synaptic proteomics, *Ddx3x*^flox/flox^ females were crossed with B6.Cg-Edil3^Tg(Sox2–Cre)1Amc^/J males (Sox2-Cre/+) ^82^ (The Jackson Laboratory, stock number #008454). To generate the *Emx1-Ddx3x* mouse line used for the behavioral testing, *Ddx3x*^flox/flox^ females were crossed with B6.129S2-*Emx1*^tm1(cre)Krj^/J (Emx1^IRES-Cre^/+) hemizygous males ^59,60^ (The Jackson Laboratory, stock number #005628). The colony was maintained in a room on a 12/12 hours light/dark cycle, with lights on at 7 A.M. at a constant temperature of 21-22°C and 55% humidity. Standard rodent chow and potable water were available *ad libitum.* Animals were socially housed, with 3-5 mice per cage. Mice were weaned at P21. The colony was maintained on a C57BL/6J background. Genotyping is described in the **Supplemental Note**.

### Primary cortical cultures

Cultures were prepared from E15 mouse embryos with a modified protocol ^83^. Cortices from single embryos were isolated in ice-cold Hank’s balanced salt solution (HBSS) and then digested with pre-warmed papain containing DNase (Worthington, #LK003150) at 37°C and 5% CO_2_ for 30 min. Tissue chops were washed 3 times with Neuronal Plating Medium and then centrifuged for 3 min at 400 g. Mechanical dissociation was then performed in 1 ml of pre-warmed Neurobasal Medium (Thermo Fisher Scientific, #21103049). Cell density was determined with the Countess Automated Cell Counter (Thermo Fisher Scientific), with a yield on the order of 3-4 × 10^6^ cells per embryo. Irrespective of timing of transfection/infection (see below), cells were maintained in Neuronal Plating Medium, and 90 min after plating the medium was replaced with pre-warmed Neurobasal Plus (Thermo Fisher Scientific, #A3582901) supplemented with B27 Plus (Thermo Fisher Scientific, #A3582801). At days in vitro (DIV) 4, cells were treated with 1 µM cytosine arabinoside (1-b-D-arabinofuranosylcytosine (Sigma-Aldrich Inc, #C1768). Genotyping of the cultures is described in the Supplemental Note.

### DNA transfections

For dendritic branching analyses, right before plating (DIV0) 2 × 10^5^ cells were incubated for 1 hour in a 50 µl of Neurobasal containing 0.5 µl of Lipofectamine (Thermo Fisher Scientific, #11668019) and 500 ng of pAAV-Ef1a-mCherry-IRES-Cre (Addgene, #55632) in 1.5 ml conical tubes and then seeded onto poly-L-lysine-coated coverslips in 24-well plates containing Neuronal Plating Medium. To assess dendritic spines, 1.5 × 10^5^ neurons were plated at DIV0 onto poly-L-lysine-coated coverslips in 24-well plates containing Neuronal Plating Medium and then transfected at DIV9 by adding 100 µl of Neurobasal containing 0.75 µl of Lipofectamine and 500 ng of pAAV-Ef1a-mCherry-IRES-Cre directly onto each well. For nucleoli analyses, cells were transfected as for dendritic branching analyses with 500 ng of pAAV-Ef1a-mCherry-IRES-Cre or 500 ng of a control pAAV-Ef1a-mCherry (Addgene, #114470).

### Adeno-associated virus transductions

At DIV0, 10^6^ cells were directly plated onto poly-L-lysine-coated 35 mm Petri dishes containing Neuronal Plating Medium, and cells were then transduced with 1 µl of AAV8-Ef1a-mCherry-IRES-Cre (Addgene, #55632-AAV8, 1X10^13^ vg/ml, multiplicity of infection (MOI) of 10,000). Cells were harvested at DIV9.

### Immunostainings on cultures

At DIV9 (dendritic branching analyses and nucleoli analyses) or DIV14 (dendritic spine analyses), coverslips were washed with pre-warmed 1X phosphate buffered saline (PBS) and fixed with a pre-warmed solution of 4% paraformaldehyde (PFA) /4% sucrose in 1X PBS at room temperature (RT) for 10 min. Coverslips were then washed once with 50mM glycine in 1X PBS and twice with 1X PBS /0.1% Triton X-100 at RT for 5 min, and then treated with 1X PBS/0.3% Triton X-100 for 20 min to allow permeabilization. After blocking in 10% donkey serum/1X PBS /0.1% Triton X-100 for 1 hour at RT, coverslips were incubated with goat anti-mCherry polyclonal antibody (SICgen antibodies, #AB0040, 1:200), a rabbit anti-DDX3X polyclonal antibody (Thermo Fisher Scientific, #A300-474A, 1:400), a rabbit anti-Fibrillarin polyclonal antibody (Abcam, #ab5821, 1:2,000), and/or Hoechst dye (Thermo Fisher Scientific, #H3570, 1:1,000), diluted in blocking solution overnight at 4°C. Coverslips were then washed 3 times with 1X PBS and incubated for 1 hour at RT with a donkey anti-rabbit antibody conjugated with Alexa Fluor™ 647 (Thermo Fisher Scientific, #A-31573, 1:200) and donkey anti-goat antibody conjugated with Alexa Fluor™ 568 (Thermo Fisher Scientific, #A-11057, 1:200) in blocking solution. Coverslips were mounted on 25 x 75 x 1.0 mm microscope glass slides, on a drop of ProLong Diamond Antifade Mountant (Thermo Fisher Scientific, #P36970).

### Immunofluorescence on tissue

E18 or P0 mouse brains were fixed in 4% PFA in 1X PBS overnight at 4°C, cryopreserved in 30% (g/vol) sucrose/0.05% sodium azide/100 mM glycine in 1X PBS, and embedded in Scigen Tissue-Plus™ O.C.T. Compound (Fisher Scientific, #23-730-571). 30 μm sections were obtained using a Leica CM1860 cryostat and mounted on slides. For CRE staining, slides were washed in 1x Tris-buffered saline (TBS) 3 times, 5 min each, and blocked for 1h at RT in 10% donkey serum (Sigma-Aldrich Inc, #D9663)/0.3% Triton-X 100 in 1X TBS. Slides were incubated overnight with a guinea pig anti-CRE polyclonal antibody (Synaptic Systems, #257004, 1:100), a rabbit anti-DDX3X polyclonal antibody (Thermo Fisher Scientific, #A300-474A, 1:400), a rabbit anti-Fibrillarin polyclonal antibody (Abcam, #ab5821, 1:2000), and/or Hoechst dye (Thermo Fisher Scientific, #H3570, 1:1,000), in incubation buffer (5% donkey serum/0.3% Triton-X 100 in 1X TBS). After washing, slides were incubated with donkey anti-rabbit antibody conjugated with Alexa Fluor™ 647 (Thermo Fisher Scientific, #A-31573, 1:200) and anti-goat anti-guinea pig conjugated with Alexa Fluor™ 488 (Thermo Fisher Scientific, ab150185, 1:200) for 1h at RT. Coverslips were mounted using Dapi-Fluoromount G (SouthernBiotech, #0100-20).

### Confocal imaging

Images were acquired using a Leica TCS SP8 inverted microscope and Leica LAS X software. Excitation beams 561 and 633 nm were coupled, image size was set at 1024 × 1024 pixels, scan speed was selected at 600 Hz. 60 z-stacks of 300 nm step size (18 µm in total) were taken using a 20X air objective for dendrite analyses, and a 100X oil immersion objective for spine analyses. Z-stacks projections were prepared using the Fiji software version 2.9.0/1.53t ^84^. For nucleoli and nuclei imaging, 30 z-stacks of 300 nm step size (9 μm in total) were taken using a 100X oil immersion objective with a zoom 2. Nucleoli and nuclei volumes were reconstructed from the Z-stacks images using IMARIS 10.1.0. Nucleoli and nuclei sizes, number and nucleoli/nuclei volumes ratio were measured.

### Sholl analysis

mCherry+ neurons were reconstructed using the Simple Neurite Tracer (SNT) plugin of the Fiji software. Dendrites were classified in primary, secondary, tertiary, and quaternary based on their outgrowth from the soma and branching patterns. Dendritic lengths were also measured. Axonal projections were excluded from the analysis. Concentric rings 10 µm apart around the somata of SNT reconstructed neurons and radiating outward were defined on SNT reconstructed neurons, and the number of intersections between the dendritic arbors and the concentric rings was counted.

### Dendritic spine analyses

Dendritic spines were annotated on mCherry+ neurons starting at 30 µm from the soma and on 10 µm-long segments on the dendrite. Spines were manually counted on an average of 16 10 µm-long segments per neuron, and spine count was then averaged per neuron. Dendritic spine density was measured as the total number of dendritic spines in 10 µm of dendritic length. For each spine, the head width (W) and neck length (L) was measured. Spines were classified into 6 morphological subtypes: branched (two-head spines), mushroom (W ≥ 0.7 µm; 1 < L > 3 µm), stubby (W ≤ 0.6 µm; no neck detected), thin (W ≤ 0.7 µm; L ≤ 1 µm), long thin (W ≤ 0.7 µm; 1 < L > 3 µm), and filopodia (no head detected; L ≥ 3 µm), based on previous literature ^18,85^.

### Synaptosomes

Synaptosomes were isolated as described previously ^18^. P21 *Sox2-Ddx3x*^+/+^ and *Sox2-Ddx3x*^+/−^ female mice were euthanized by cervical dislocation. Cortices were rapidly dissected on ice and homogenized in ice-cold homogenizing buffer (0.32M sucrose, 1mM EDTA, 1mg/ml BSA, 5mM HEPES pH=7.4) in a glass Teflon® douncer, and centrifuged at 3,000 g for 10 min at 4°C. The supernatants were recovered and centrifuged at 14,000 g for 12 min at 4°C. The pellets containing synaptosomes were gently resuspended in Krebs-Ringer buffer (140mM NaCl, 5mM KCl, 5mM glucose, 1mM EDTA, 10mM HEPES pH=7.4), subjected to a density gradient upon addition of Percoll™ Plus (Sigma-Aldrich) (final 45% v/v), and centrifuged at 14,000 rpm for 2 min at 4°C to enrich the synaptosomes at the surface of the gradient. The synaptosomes were recovered, washed in Krebs Ringer buffer, and then centrifuged at 14,000 rpm for 30 sec at 4°C. The pellets containing purified synaptosomes were lysed as detailed in the proteomics section.

### Sample preparation for liquid chromatography with tandem mass spectrometry (LC-MS/MS)

Cell or synaptosome pellets were suspended in 200 μl RIPA buffer contains protease and phosphatase inhibitor cocktail. Three bursts (5% amp, 1 second pulses) of sonication for 15 sec each were used to lyse the cells/synaptosomes. Suspension was centrifuged at 14,600 rpm for 10 min at 4°C, and 175 μl of the supernatant was aliquoted for chloroform:methanol:water protein precipitation using standard protocols. The dried protein pellet was resuspended in 10 μl 8M urea containing 400 mM ammonium bicarbonate (ABC), reduced with dithiothreitol (DTT), alkylated with iodoacetamide, and dual enzymatic digestion with LysC and trypsin (carried out at 37°C overnight or 7 hrs, respectively). Digestion was quenched by adding trifluoroacetic acid (to 0.5%) prior to peptide de-salting with C18 MiniSpin columns (The Nest Group). The effluents from the de-salting step were dried and re-dissolved in in MS loading buffer (2% acetonitrile, 0.2% trifluoroacetic acid). Protein concentrations (A260/A280) were measured with a Nanodrop 2000 UV-Vis Spectrophotometer (Thermo Fisher Scientific). Each sample was then further diluted with MS loading buffer to 0.06µg/µl, with 300ng (5µl) injected for LC-MS/MS analysis. 1:10 dilution of 10X Pierce Retention Time Calibration Mixture (#88321) was added to each sample to check for retention time variability during LFQ data analysis.

### LC-MS/MS analyses

LC-MS/MS analysis was performed on a Thermo Scientific Q Exactive HF-X equipped with a Waters M-class UPLC system utilizing a binary solvent system (A: 100% water, 0.1% formic acid; B: 100% acetonitrile, 0.1% formic acid). Trapping was performed at 10 µl/min, 99% Buffer A for 3 min using a Waters nanoEase M/Z Symmetry C18 Trap Column (100A, 5 µm, 180 µm x 20mm). Peptides were separated using a nanoEase M/Z Peptide BEH C18 Column (130Å, 1.7 µm, 75 µm X 250 mm) (37°C) by elution with linear gradients reaching 6% B at 2 min, 25% B at 175 min, 40% B at 195 min, and 90% B at 200 min. Column regeneration and up to three blank injections were carried out in between all sample injections. MS was acquired in profile mode over the 350-1,500 m/z range using 1 microscan, 60,000 resolution, AGC target of 3E6, and a maximum injection time of 100 ms. Data dependent MS/MS were acquired in centroid mode on the top 20 precursors per MS scan using 1 microscan, 30,000 resolution, AGC target of 1E5, maximum injection time of 100 ms, and an isolation window of 1.6 m/z. Precursors were fragmented by HCD activation with a collision energy of 28%. MS/MS were collected on species with an intensity threshold of 2.5E4, charge states 2-6, and peptide match preferred. Dynamic exclusion was set to 20 sec. The LC-MS/MS data were processed with Progenesis QI software (Nonlinear Dynamics, version 4.2) with protein identification carried out using in-house Mascot search engine (2.4). The Progenesis QI software performs chromatographic/spectral alignment (one run is chosen as a reference for alignment of all other data files to), mass spectral peak picking and filtering (ion signal must satisfy the 3 times standard deviation of the noise), and quantitation of proteins and peptides. A normalization factor for each run was calculated to account for differences in sample load between injections as well as differences in ionization. The normalization factor was determined by calculating a quantitative abundance ratio between the reference run and the run being normalized, with the assumption being that most proteins/peptides are not changing in the experiment so the quantitative value should equal 1. The experimental design was setup to group multiple injections (technical and biological replicates) from each run into each comparison sets. The algorithm then calculates the tabulated raw and normalized abundances, ANOVA *P*-values for each feature in the data set. The MS/MS spectra was exported as mgf (Mascot generic files) for database searching. Mascot Distiller was used to generate peak lists, and the Mascot search algorithm was used for searching against the Swiss Protein database with taxonomy restricted to *Mus musculus* (17,174 sequences); and carbamidomethyl (Cys), oxidation of Met, Phospho (Ser, Thr, Tyr), acetylation (Lys and Protein N-term), and deamidation (Asn and Asp) were entered as variable modifications. Two missed tryptic cleavages were allowed, precursor mass tolerance was set to 10 ppm, and fragment mass tolerance was set to 0.02 Da. The significance threshold was set based on a False Discovery Rate (FDR) of 2%. The Mascot search results was exported as .xml files and then imported into the processed dataset in Progenesis QI software where peptides identified were synched with the corresponding quantified features and their corresponding abundances. Protein abundances (requiring at least 1 unique peptide with MOWSE score of >95% confidence) were then calculated from the sum of all non-conflicting peptide ion ID assignment for a specific protein on each run. A label free quantitation approach utilizing Progenesis QI (Nonlinear Dynamics)^86^ was utilized to obtain quantitative information on peptides and proteins.

### Developmental milestones

Testing was performed as reported previously ^22^, using a battery of tests adapted from the Fox scale ^87,88^. Details are described in the **Supplemental Note.**

### Gait analyses

Details are described in the **Supplemental Note.**

### Adult motor testing

Testing started at 10 weeks +/- one week. Behavioral testing was conducted during the light phase in sound-attenuated rooms. Mice were handled daily for one week prior to behavioral testing to assess general health, physical appearance, and spontaneous activity (**Supplemental Note**). Mice were habituated to the testing room for 30 min prior to testing. All surfaces and equipment were cleaned with Rescue Veterinary disinfectant between trials and nitrile gloves were used. The four genotypes were tested on the same day in randomized order by an experimenter blind to genotype. All data were scored and analyzed blind to the genotype. For accelerating rotarod, mice were tested on a motorized rod measuring 3 cm in diameter (Omnitech Electronics Inc) with gradual speed increase from 4 to 40 rpm over 5 min. Mice were presented with four 5 min-trials per day, 1 hour apart, over three consecutive days. The latency to fall and speed was recorded. The balance beam test was conducted as described previously ^22^. Briefly, the beam consisted of a 70-cm long square prism with 1.3 cm face placed horizontally 40 cm above the bench. Each frame was 5-cm long. Mice were allowed to freely walk on the beam for two sessions (one training and one testing) of 2 min each, separated by 4 hrs. The number of slips and the distance covered during the testing session were analyzed. The vertical pole test was conducted as described previously ^22^. A wooden pole wrapped in tape to facilitate walking was used. The test consisted of two consecutive training days and a testing day (3 trials each day). Mice were placed facing upwards just below the top of the pole and the time to complete the turn and the time to climb down the pole were measured. The wire hanging test was conducted as described previously ^22^. Mice were led from the base of the tail and allowed to grasp a rod placed 40 cm above the bench by both forepaws. The task consisted of a 5 sec-training session followed by three 60 sec-testing sessions. The latency to fall and the distance traveled along the hanger were recorded, and the average of the testing sessions was used for analysis.

### Statistical analyses

All statistics and plots were generated using custom R scripts. Statistical tests, number of animals in each experiment, and significance are indicated in each figure legend. Additional statistics are indicated in figure legends. Data are shown as mean ± SEM. Outliers are defined as data points below Q1-1.5xIQR or above Q3+1.5xIQR, where Q1 is the first quartile, Q3 is the third quartile, and the IQR is the interquartile range. Outliers, shown in plots with the ≅ symbol, were removed from the calculations of the mean and SEM and for the statistical tests. Repeated measure ANOVA was used for repeated observations. For comparisons between two groups, Shapiro–Wilk test was used to assess normality, followed by Student’s t-test for normally distributed data with equal variance, Welch Two Sample t-test for normally distributed data with unequal variance, or Wilcoxon signed-rank test for not normally distributed data. In case of multiple comparisons, two-way ANOVA test was employed. To compare probability distributions, the Kolmogorov–Smirnov test was used. Whenever possible, data are reported by individual (not only by group).

## ACKNOWLEDGMENTS

Research reported in this publication was supported by the Eunice Kennedy Shriver National Institute Of Child Health & Human Development of the National Institutes of Health under Award Number R21HD097561 and R01 R01HD104609 (to SDR). The content is solely the responsibility of the authors and does not necessarily represent the official views of the National Institutes of Health. The work was also supported by the Beatrice and Samuel A. Seaver Foundation and a Distinguished Scholar Award from the Icahn School of Medicine at Mount Sinai to SDR. AM, LD, and JL were supported by fellowships from the Beatrice and Samuel A. Seaver Foundation. AM was also supported by the Friedman Brain Institute (Doft Family and Friedman Brain Institute Postdoctoral Innovator Award) and the 2023 Brain and Behavior Research Foundation Young Investigator Grant. LD was also supported by a traineeship from the NIDCR-Interdisciplinary Training in Systems and Developmental Biology and Birth Defects (NICHD, T32HD075735). ZA was supported by a Dean’s Undergraduate Research Fund Grant from New York University and Seaver Undergraduate Research Scholarship. We also thank the Keck MS & Proteomics Resource at Yale School of Medicine for providing the necessary mass spectrometers and the accompany biotechnology tools funded in part by the Yale School of Medicine and by the Office of The Director, National Institutes of Health (S10OD02365101A1, S10OD019967, and S10OD018034). The funders had no role in study design, data collection and analysis, decision to publish, or preparation of the manuscript. We thank Dévina Ung for initial experimental setup, Kristi Niblo for technical support, and Olivia Pistone, Sylvia Maxwell and Mahmuda Hannan for support with data entry.

## AUTHOR CONTRIBUTIONS

SDR, AM, LD, JL conceptualize the study and designed the experiments. AM, LD, JL, YP, CF, ZA, and MGF conducted experiments and collected data. SDR, AM, LD, JL analyzed the data. SDR, AM, LD, JL wrote the original draft and all the other authors edited and approved the manuscript.

## COMPETING INTERESTS

The authors declare no competing interests.

## RESOURCE AVAILABILITY

Further information and requests for resources and reagents should be directed to and will be fulfilled by the

Lead Contact, Silvia De Rubeis.

## Notes

### Competing Interest Statement

The authors have declared no competing interest.

